# Atypical genomic patterning of the cerebral cortex in autism with poor early language outcome

**DOI:** 10.1101/2020.08.18.253443

**Authors:** Michael V. Lombardo, Lisa Eyler, Tiziano Pramparo, Vahid H. Gazestani, Donald J. Hagler, Chi-Hua Chen, Anders M. Dale, Jakob Seidlitz, Richard A. I. Bethlehem, Natasha Bertelsen, Cynthia Carter Barnes, Linda Lopez, Kathleen Campbell, Nathan E. Lewis, Karen Pierce, Eric Courchesne

## Abstract

Cortical regional identities develop through anterior-posterior (A-P) and dorsal-ventral (D-V) prenatal genomic patterning gradients. Here we find that A-P and D-V genomic patterning of cortical surface area (SA) and thickness (CT) is intact in typically developing and autistic toddlers with good language outcome, but is absent in autistic toddlers with poor early language outcome. Genes driving this effect are prominent in midgestational A-P and D-V gene expression gradients and prenatal cell types driving SA and CT variation (e.g., progenitor cells versus excitatory neurons). These genes are also important for vocal learning, human-specific evolution, and prenatal co-expression networks enriched for high-penetrance autism risk genes. Autism with poor early language outcome may be linked to atypical genomic cortical patterning starting in prenatal periods and which impacts later development of regional functional specialization and circuit formation.

**One Sentence Summary:** Genomic patterning of the cortex is atypical in autistic toddlers with poor early language outcome.

---

It is widely accepted that the autisms (ASD) arise in large part due to complex genetic mechanisms (*1*, *2*). The major priorities for the field are to develop an individualized understanding of how complex genetic mechanisms cascade over development to cause phenotypic differentiation at multiple scales and link these mechanisms to clinical outcomes with high real-world impact and relevance (*3–5*). At the nexus of these priorities, our prior functional imaging (fMRI) work showed that large-scale activity in blood leukocyte gene co-expression modules differentially relates to language-relevant functional neural phenotypes measured in typically developing (TD) and ASD toddlers with good (ASD Good) versus poor (ASD Poor) early language outcome (*6*). This result indicates that the atypical language-relevant functional neural phenotypes typically seen in ASD Poor (*7*) are driven by different underlying functional genomic mechanisms.

The genes of importance that differentially relate to language-relevant functional neural phenotypes in ASD Poor (*6*) are an omnigenic (*8*) array of genes that are typically broadly expressed across many tissues including the brain. Broadly expressed genes tend to be one of the most important classes of ASD-risk genes and in the brain they show peak expression during early prenatal periods, when cell proliferation, differentiation, neurogenesis, and migration are the primary biological processes (*9, 10*). If these genes operate at early prenatal periods to affect proliferation, differentiation, neurogenesis, and migration, this suggests that structural features of the developing cerebral cortex such as surface area (SA) and cortical thickness (CT) may be substantially altered in the ASD Poor subtype.

The action of broadly expressed genes during prenatal periods may also be important for how the cortex is genomically patterned. It is well established that during prenatal periods the cortex is patterned by gene expression gradients that follow anterior-posterior (A-P) and dorsal-ventral (D-V) axes (*11–17*). This prenatal genomic patterning is the beginning of cortical arealization processes that allow different cortical regions to develop their own cellular, functional, and circuit identities (*11*, *13*, *15*, *16*). Cortical arealization or patterning may be atypical in ASD. Prior evidence from case-control comparisons of post-mortem cortical tissue has found dysregulation of cortical patterning genes and attenuation of gene expression differences in frontal versus temporal cortex (*18–20*). WNT-signaling is known to affect cortical patterning (*13*, *15*, *16*, *21*) and WNT-signaling abnormalities are also identified in ASD (*19*, *20*, *22*–*24*), particularly within broadly expressed ASD-risk genes (*10*). Therefore, if broadly expressed genes in early prenatal periods impact the ASD Poor subtype, could this also implicate a disturbance of prenatal genomic patterning of the cortex?

In the current work we examined these questions in a sample of n=123 toddlers (12-50 months) with and without ASD (ASD Good n=38, ASD Poor = n=38, and TD n = 47). With T1-weighted structural MRI images, we used Freesurfer (http://surfer.nmr.mgh.harvard.edu) to extract SA and CT measures from 12 cortical regions that parcellate the cortex by hierarchical genetic similarity (*25–27*). This cortical parcellation, known as GCLUST, was chosen in order to maximize sensitivity for detecting genetic relationships (*28*). GCLUST is also sensitive to the SA and CT genetic similarity gradients that fall along A-P and D-V axes and therefore, also maximizes sensitivity for detecting such genetically sensitive A-P and D-V gradient effects (*25–27*) (see Methods for more details).

Since one of the most robust findings on early structural brain development in ASD is the on-average effect of early brain overgrowth in the first years of life (*4*, *29*–*31*), we started by examining whether there are subtype differences on global measures such as total cortical volume (CV), SA and mean CT. Statistical models controlling for age and sex identified a group effect on total CV (*F(2,193)* = 14.30, *p* = 2.74e-6, *η*^*2*^ = 0.075) that is driven by the ASD Poor subtype having on-average larger CV than the other groups (ASD Good vs ASD Poor *t(125)* = 1.88, *p* = 0.06, *Cohen’s d* = −0.42; TD vs ASD Poor *t(132)* = −2.98, *p* = 0.003, *Cohen’s d* = −0.70; TD vs ASD Good *t(127)* = −1.10, *p* = 0.27, *Cohen’s d* = −0.26) (Fig. 1A). A group effect also emerged for total SA (*F(2,193)* = 15.39, *p* = 1.14e-6, *η*^*2*^ = 0.072) and was again driven by on-average increases in ASD Poor relative to the other groups (ASD Good vs ASD Poor *t(125)* = 1.79, *p* = 0.07, *Cohen’s d* = −0.40; TD vs ASD Poor *t(132)* = −2.84, *p* = 0.005, *Cohen’s d* = −0.71; TD vs ASD Good *t(127)* = −1.28, *p* = 0.20, *Cohen’s d* = −0.27) (Fig. 1B). In contrast, no group differences were identified for mean CT (*F(2,193)* = 2.80, *p* = 0.06, *η*^*2*^ = 0.002; ASD Good vs ASD Poor *t(125)* = −0.24, *p* = 0.80, *Cohen’s d* = 0.03; TD vs ASD Poor *t(132)* = 0.17, *p* = 0.86, *Cohen’s d* = 0.11; TD vs ASD Good *t(127)* = 0.37, *p* = 0.71, *Cohen’s d* = 0.07). These effects illustrate that the ASD Poor subtype drives the on-average effect of early brain overgrowth in autism. Differences in CV and total SA, but not mean CT, is compatible with other work showing that early brain overgrowth is largely driven by expansion of cortical SA rather than CT (*32, 33*). We next examined regional level SA or CT effects when adjusting for global differences using the GCLUST parcellation (see Methods). Here we find no evidence of SA or CT group differences for any of the 12 GCLUST regions, indicating that the primary overall group differences in brain size are restricted to global effects in CV and SA, rather than localized regional effects after adjusting for such global effects.

**Fig. 1:**
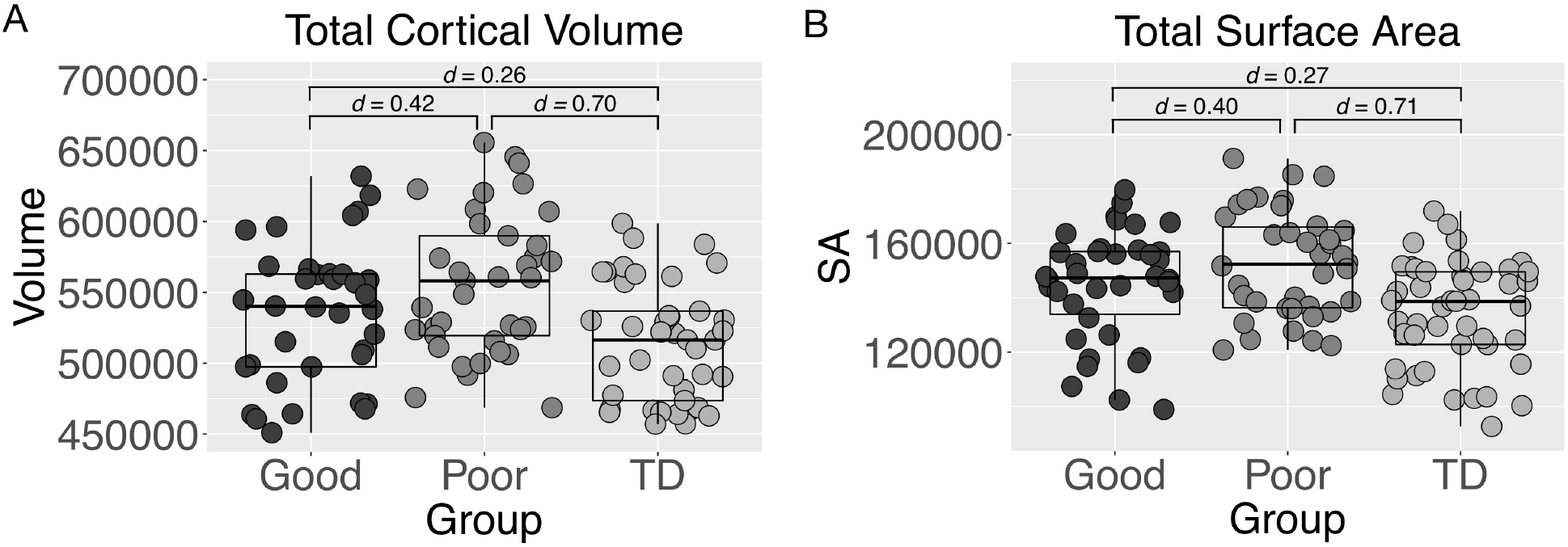
Subtype differences in total cortical volume (A) and total surface area (B). Standardized effect sizes (Cohen’s d) are shown for each pairwise group comparison.

We next examined large-scale associations between gene expression and regional SA or CT from the GCLUST parcellation. To examine gene expression, leukocyte cells were extracted from blood samples and microarrays were used to quantify expression from 14,426 protein coding genes. This set 14,426 genes was then reduced to 21 gene co-expression modules using weighted gene co-expression network analysis (WGCNA) (*34*). We then we used partial least squares (PLS) analysis to test for large-scale associations between blood leukocyte co-expression modules and SA or CT phenotypes from the GCLUST parcellation (see Methods for more details). For SA, we identified one statistically significant latent variable (LV) pair (SA LV1: d = 3.99, p = 0.0001), which explains 36% of the covariance between SA and gene expression. To decompose how this multivariate relationship manifests across co-expression modules and groups, in Fig. 2D we show which co-expression modules have ‘non-zero’ relationships in each group. These ‘non-zero modules’ have 95% confidence intervals (CIs) estimated by bootstrapping that do not include a correlation of 0 and are thus the most important co-expression modules driving the SA LV1 relationship. In contrast, co-expression modules that we dub as ‘zero modules’ are those whereby the 95% CIs include a correlation of 0 and thus do not reliably contribute to the overall SA LV1 relationship. Non-zero modules for SA LV1 account for a good majority (68%) of all genes examined and this effect is compatible with ideas about omnigenic effects on complex traits such as imaging phenotypes in ASD subtypes (*6, 8*). Fig. 2D also shows that non-zero modules are highly similar for ASD Good and TD groups, whereas hardly any non-zero modules are present for ASD Poor. This similarity between ASD Good and TD can be quantified as a significant positive correlation in the PLS correlations values for these groups (Fig. 2D-E) (r = 0.55, p = 0.008). This result indicates that the SA LV1 relationship manifests similarly in TD and ASD Good groups. In contrast, there is a lack of correlation between ASD Poor and the other groups (ASD Poor-ASD Good: r = 0.27, p = 0.23; ASD Poor-TD: r = −0.41, p = 0.06). Therefore, SA LV1 can be described as a large-scale SA-gene expression relationship that likely reflects a normative phenomenon present in TD and which is also preserved in the ASD Good subtype. However, this normative SA-gene expression relationship is absent in the ASD Poor subtype.

**Fig. 2:**
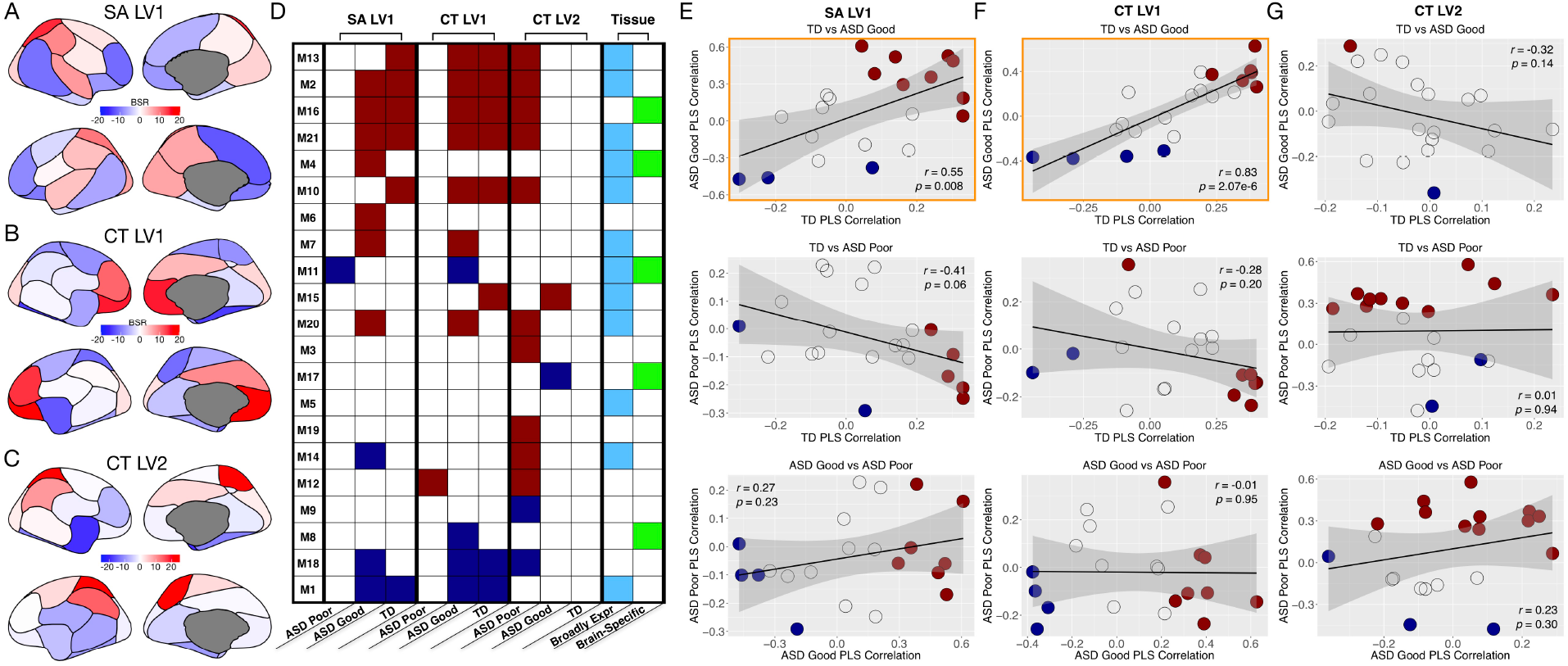
Multivariate gene co-expression relationships with SA and CT. Panels A-C show brain bootstrap ratios (BSR) for SA LV1 (A), CT LV1 (B), and CT LV2 (C) for all 12 regions from the GCLUST SA and CT parcellations. Regions that are increasingly colored red and blue are regions that most reliably contribute to the PLS relationship. Panel D shows which co-expression modules are ‘non-zero’ modules (dark red or dark blue) or ‘zero’ modules (white). Non-zero modules are co-expression modules where the correlation between gene expression and SA or CT is significantly non-zero, as indicated by 95% bootstrap confidence intervals not encompassing a correlation of 0. These non-zero modules are the strongest contributors to the PLS relationship. All white cells indicate ‘zero’ modules that are not sufficiently correlated in a non-zero way (e.g., 95% bootstrap confidence intervals include a correlation of 0). Non-zero modules in dark red can be interpreted as positive correlations with brain regions in panels A-C colored in red. However, for brain regions colored in blue, the correlations in non-zero modules colored in dark red are interpreted as negative correlations. These interpretations about the directionality of the correlation are reversed when it comes to non-zero modules colored in dark blue. The final two columns show which modules are enriched for broadly expressed or brain-specific genes. Panels E-G show similarity in PLS correlations for all pairwise comparisons for SA LV1 (E), CT LV1 (F), and CT LV2 (G). In these scatterplots each dot is a co-expression module and the x and y-axes indicate the PLS correlations for different groups. Dots colored in dark red and dark blue indicate the non-zero modules, while grey dots indicate zero modules. Scatterplots with the orange outline indicate similar relationships for TD and ASD Good for SA LV1 and CT LV1.

PLS analysis applied to CT data isolated 2 statistically significant LV pairs (CT LV1: d = 4.30, p = 0.0001; CT LV2: d = 3.09, p = 0.0001), explaining 37% and 19% of the covariance between CT and gene expression respectively. Similar to SA LV1, non-zero modules for CT LV1 comprise a large majority of all genes examined (65%) and are highly similar for ASD Good and TD, but not ASD Poor (Fig. 2D, F, G). These results indicate that CT LV1 mostly pertains to a normative relationship preserved across TD and ASD Good, but which is absent in ASD Poor. In contrast to CT LV1, the non-zero modules for CT LV2 are almost exclusively relevant for the ASD Poor subtype, comprise about 48% of all genes examined, and do not show strong correlations between groups (Fig. 2D, F, G). These results indicate that CT LV2 captures a relationship that is specific to ASD Poor.

Given that our prior work discovered that PLS non-zero modules related to language-relevant functional neural phenotypes are highly enriched for broadly expressed genes (*6*), we next asked if SA and CT non-zero modules were similarly enriched. Indeed, SA LV1 non-zero modules are highly enriched in broadly expressed genes (enrichment odds ratio (OR)= 3.48, p = 1.90e-71) but not brain-specific genes (OR = 1.67, p = 0.23), while no enrichments were present for zero modules (broadly expressed, OR = 1.10, p = 0.99; brain-specific, OR = 0.94, p = 0.99). CT LV1 non-zero modules are also highly enriched in broadly expressed genes (OR= 2.96, p = 4.43e-43) but not brain-specific genes (OR = 1.56, p = 0.55), while zero modules were not enriched in either broadly expressed (OR = 1.10, p = 0.99) or brain-specific genes (OR = 0.94, p = 0.99). In contrast, CT LV2 showed enrichments for broadly expressed genes in both non-zero (OR= 1.90, p = 1.31e-7) and zero modules (OR = 2.43, p = 1.34e-28), but no enrichments for brain-specific genes (non-zero modules OR = 1.21, p=0.98; zero modules OR = 1.40, p = 0.57). These results show that SA and CT LV1 results are largely driven by the class broadly expressed genes, while for CT LV2 the enrichment for broadly expressed genes is present, but not specific to non-zero modules.

We next investigated how genomic variability patterns SA and CT cortical phenotypes. The patterning of PLS brain bootstrap ratios (BSR) shown in Fig. 2A-C can be used to answer this question. BSRs indicate the directionality through which gene expression is associated with SA and CT and can also show how these relationships manifest similarly or differently across brain regions. It is visually evident from Fig. 2A-C that BSR patterning is not uniform across cortical regions and varies considerably along A-P and D-V axes. With a 2-cluster solution previously identified by Chen and colleagues (*25–27*) to be the genetically parcellated A-P and D-V axes of SA and CT (Fig. 3A), we confirm that BSRs highly differ along these A-P and D-V clusters (Fig. 3C, E, G). This indicates that the relationship between gene expression and SA or CT at one pole of the A-P or D-V axes is different relative to the other pole.

**Fig. 3:**
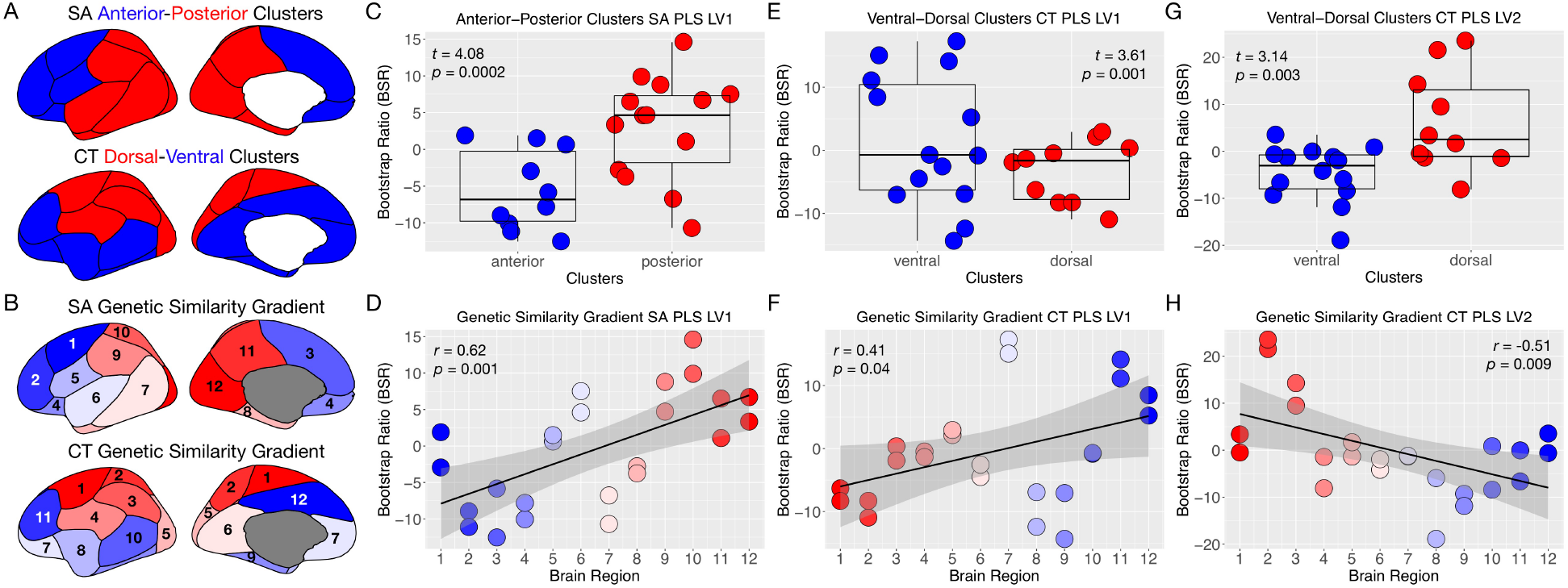
Cortical patterning along genetic similarity gradients. Panel A shows the coarse 2-cluster anterior-posterior (A-P) and dorsal-ventral (D-V) genetic similarity partitions identified by Chen and colleagues (25–27). Panel B shows the rank ordering of regions by hierarchical genetic similarity gradients discovered by Chen and colleagues (25–27). These two parcellations were utilized to examine how brain BSRs may vary along these genetic similarity gradients. Panels C-D show A-P and genetic similarity gradients for SA LV1. Panels E-H show D-V (E, G) and genetic similarity gradients (F, H) for CT LV1 (E, F) and CT LV2 (G, H).

Perhaps even more striking than these differences between binary A-P and D-V partitions is that BSRs also covary along continuous A-P and D-V genetic similarity gradients. After ordering regions by genetic similarity gradients discovered by Chen and colleagues (*25–27*) (Fig. 3B) we find that BSRs are highly correlated with the ordering along this axis of genetic similarity between regions (Fig. 3D, F, H). This indicates that large-scale blood leukocyte gene co-expression relationships with SA and CT reveal how the cortex is genomically patterned to promote the development of cortical regionalization and areal identity (*13*). Because SA LV1 and CT LV1 are normative effects primarily relevant for TD and ASD Good, but not ASD Poor, these results indicate that normative genomic patterning of the cortex does not occur in the ASD Poor subtype. Conversely, CT in ASD Poor subtype may be patterned in a completely different way given that CT LV2 was primarily relevant to this subtype and given that the BSR patterning is reversed for CT LV2 compared to CT LV1 (Fig. 3E and G versus Fig. 3F and H). Given evidence of focal laminar patches throughout the cortex in ASD (*35*), it will be important for future work to investigate further how such phenomena may be relevant to atypical CT patterning, particularly in the ASD Poor subtype.

In contrast to these effects of genomic patterning along A-P and D-V gradients, we also examined if the effect size of SA or CT difference between ASD subtypes and TD would similarly follow A-P and D-V gradients. Prior work using lobar parcellations has suggested that case-control differences in cortical size may follow an A-P gradient (*36*). However, effect sizes do not seem to follow either the 2-cluster A-P and D-V partitions or continuous genetic similarity gradients (Fig. S1). This result suggests that these cortical patterning effects are not simply effects that can be seen as on-average group differences in SA or CT and point more towards the specific importance of how the underlying genomic mechanisms act to pattern SA and CT across the cortex.

Because cortical regionalization begins in early prenatal periods from A-P and D-V gradient patterning of gene expression (*12*–*14*, *17*), we next assessed whether genes from SA and CT non-zero modules are the same genes that play important prenatal roles in the genomic gradient patterning of the cortex. Using the Development PsychENCODE dataset, we used sparse PCA (*37*) to identify A-P (PC1) and D-V (PC2) gene expression gradients and the most important genes contributing to those gradients from 12 regions of prenatal cortical tissue sampled from 12-24 weeks post-conception (e.g., midgestation) (Fig. 4A-C). Remarkably, non-zero module SA LV1 and CT LV1 gene sets are highly enriched for genes that drive the prenatal A-P and D-V gradients (Fig. 4D). CT LV2 genes were also enriched for A-P and D-V prenatal gradients, but unlike SA LV1 and CT LV1, the enrichments were apparent for both zero and non-zero modules (Fig. 4D). These results suggest that the genes responsible for the normative SA LV1 and CT LV1 relationships are also genes in prenatal periods that act to initialize the regionalization and patterning of cortex along A-P and D-V axes. Since SA LV1 and CT LV1 relationships are largely absent in the ASD Poor subtype, this result suggests that the atypical genomic patterning of SA and CT in this subtype could stem from perturbations in earlier prenatal development.

**Fig. 4:**
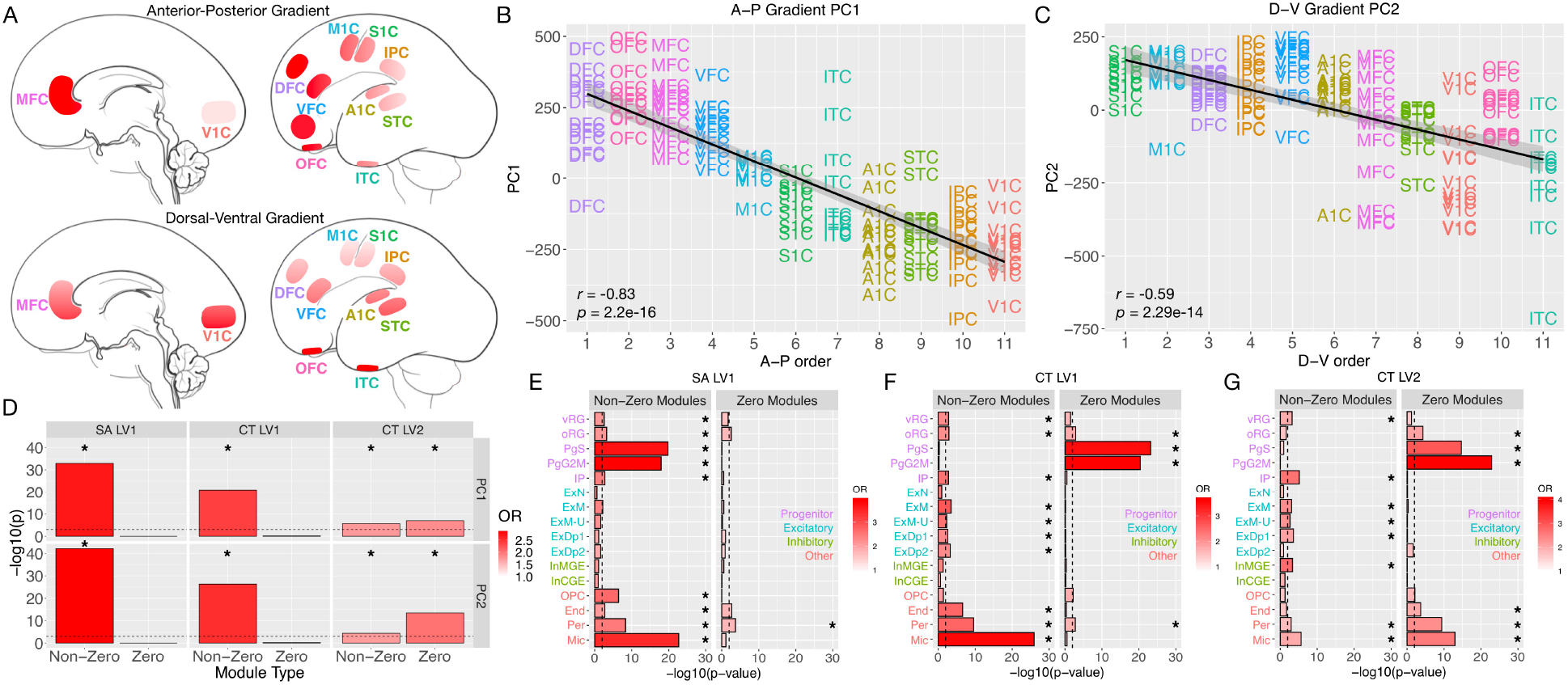
Enrichment between PLS non-zero modules and genes involved in prenatal A-P and D-V expression gradients and prenatal cell types. Panels A shows cortical brain areas sampled from 12-24 weeks post-conception from the Development PsychENCODE RNA-seq dataset from Li and colleagues (14). AC-PCA (37) was utilized to isolate anterior-posterior (A-P) (PC1, panel B) and dorsal-ventral (D-V) (PC2, panel C) expression gradients. Panel D shows −log10 p-values for enrichment tests of non-zero and zero modules for SA LV1, CT LV1, and CT LV2 for genes isolated from PC1 and PC2. Panels E-F show enrichments in prenatal cell types for SA LV1 (E), CT LV1 (F), and CT LV2 (G). Abbreviations: A-P, anterior-posterior; D-V dorsal-ventral; PC, principal component; OR, enrichment odds ratio; vRG, ventricular radial glia; oRG, outer radial glia; PgS, cycling progenitors (S phase); PgG2M, cycling progenitors (G2/M phase); IP, intermediate progenitors; ExM, maturing excitatory; ExN, migrating excitatory; ExM-U, maturing excitatory upper enriched; ExDp1, excitatory deep layer 1; ExDp2, excitatory deep layer 2; InCGE, interneuron caudal ganglion eminence; InMGE, interneuron medial ganglion eminence; OPC, oligodendrocyte precursor cells; End, endothelial cells; Per, pericytes; Mic, microglia.

The evidence that SA and CT non-zero modules are enriched for genes that are important for midgestational A-P and D-V expression gradients leaves open the question of what prenatal cell types might drive such effects. The radial unit hypothesis (*12*) suggests that symmetric cell division in progenitor cell types (e.g., radial glia) in the ventricular zone leads to a substantial proliferation of radial units that then each become their own cortical column and thus, leads to substantial expansion of SA. Variation in this proliferative process in different parts of the ventricular zone protomap regulates regional differences in SA (*12*, *13*, *38*). Programmed cell death could also be another mechanism regulating SA (*13*) and could implicate microglia involvement. In contrast, CT is likely regulated by asymmetric cell division leading to more neurons within particular cortical columns (*12*) as well as intermediate progenitor cell types (*13*). CT is also heavily influenced by dendritic arborization (*39*). While arborization changes over development due to a variety of factors such as experience-dependent pruning, CT and the trajectory it follows over development is also known to be heavily influenced by genetic factors even in middle-aged adults, suggesting that individual differences in CT have a genetic and neurodevelopmental origin (*40, 41*). Given that cell type markers from midgestational periods are available (*42*), we next asked if specific prenatal cell type markers are enriched for genes from SA and CT non-zero modules. In striking agreement with prenatal mechanisms hypothesized to affect SA expansion (*12, 13*), we find that SA LV1 non-zero modules show enrichments for all progenitor cells types - ventricular and outer radial glia (vRG, oRG), cycling progenitors in S and G2M phases of cell cycle (PgS, PgG2M), and intermediate progenitors (IP). In contrast, SA LV1 non-zero modules are devoid of enrichments in later differentiated excitatory (ExM, ExN, ExM-U, ExDp1, ExDp2) and inhibitory (InCGE, InMGE) neurons. Several non-neuronal cells also show SA LV1 enrichments, including oligodendrocyte precursors (OPC), endothelial cells (End), and microglia (Mic) (Fig. 4E; Table S1). Similar to SA LV1, CT LV1 and LV2 share enrichments for vRG progenitors. IP cell types are the only other progenitor cell type enriched for CT LV1 and CT LV2 non-zero modules, and this effect is compatible with hypothesized effects of IP cells on CT (*13*). However, CT LV1 and LV2 are differentiated from the enrichment profile of SA LV1 by the presence of enrichments with several types of excitatory neurons (Fig. 4F-G; Table S1). This result indicates a striking contrast between the SA LV1 enrichment profile of primarily progenitor cell types and are compatible with the radial unit and protomap hypotheses (*12*), differential SA and CT GWAS enrichments (*40*), and other viewpoints regarding contributors to CT (*39*). These results also highlight effects of non-neuronal cell types such as microglia cells. Microglia enrichments are present and particularly strong for SA LV1 and CT LV1 non-zero modules. This effect may have implications for programmed cell death and pruning explanations (*43*) and which may be relevant to ideas behind ASD-relevant broadly expressed genes and their particularly strong effects on non-neuronal cell types such as microglia (*10*).

The results so far suggest that SA and CT non-zero modules are highly prenatally relevant for establishing cortical patterning and regionalization and implicate several cell types that may be of mechanistic importance to different ASD early language outcome subtypes. However, are the SA and CT non-zero modules also functionally relevant for processes that are essential for language development? Our prior work showed that PLS non-zero modules associated with speech-related fMRI response (*6*) were highly enriched for differentially expressed genes in Area X from a songbird model of vocal learning (*44*). To test if similar enrichments held up for SA and CT non-zero modules we ran enrichment tests with vocal learning DE genes from Hilliard and colleagues (*44*). Remarkably, we find similar types of enrichments between DE songbird vocal learning genes and PLS non-zero modules in SA LV1 (OR = 2.02, p = 1.05e-4) and CT LV1 (OR = 1.90, p = 9.61e-4), but not zero modules (p >0.08) (Fig. 5A-C; Table S1). For CT LV2, enrichments were present at FDR q<0.05 (but not FDR q<0.01) for both non-zero (OR = 1.62, p = 0.006) and zero modules (OR = 1.61, p = 0.017). These effects suggest that many genes responsible for vocal learning in songbirds are conserved and highly represented specifically within SA and CT non-zero modules that are relevant for groups with relatively intact language (e.g., TD and ASD Good).

**Fig. 5:**
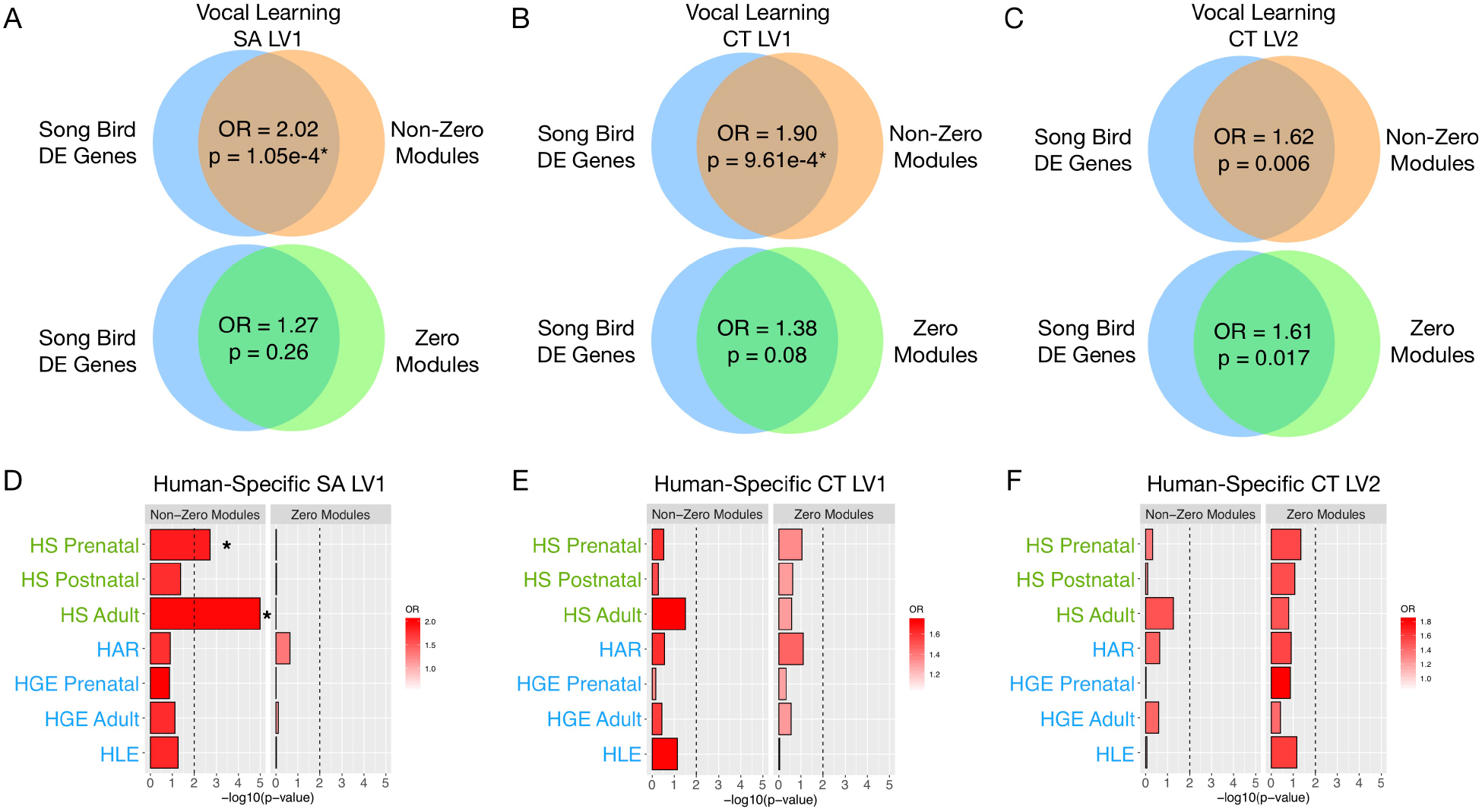
Enrichments between PLS non-zero modules and songbird vocal learning or human-specific genes. Panels A-C indicate enrichments between differentially expressed songbird vocal learning genes and non-zero and zero modules for SA LV1 (A), CT LV1 (B), AND CT LV2 (C). Panels D-F indicate enrichments between human-specific genes and non-zero and zero modules for SA LV1 (D), CT LV1 (E), and CT LV2 (F). Asterisks marks enrichments at FDR q<0.01. Abbreviations: DE, differentially expressed; OR, enrichment odds ratio; SA, surface area; CT, cortical thickness; LV, latent variable pair; HS, human-specific; HAR, human-accelerated region; HGE, human-gained enhancer; HLE, human-lossed enhancer.

Language is a uniquely human ability and there is some evidence that genes implicated in human-specific evolution are also relevant for autism (*45–48*). In prior work we found that PLS non-zero modules associated with speech-related fMRI response (*6*) were enriched for differentially expressed genes in the cortex of humans versus non-human primates (i.e. ‘human-specific’ genes). Given that cortical SA is a phenotype that is dramatically expanded in human evolution, and much moreso than CT, we investigated the hypothesis of whether SA non-zero modules would be specifically enriched for human-specific genes. Using 3 lists of human differentially expressed genes in prenatal, early postnatal, and adulthood periods (*47*), we find that SA LV1 non-zero modules are specifically enriched for prenatal and adulthood human-specific genes (prenatal OR = 1.86, p = 1.93e-3; adulthood OR = 1.97, p = 1.02e-5) (Fig. 5D; Table S1). In contrast, no such enrichments are found with genes relevant to CT LV1 or LV2 (Fig. 5E-F). In addition to differentially expressed genes we also examined genes that are targets of human-accelerated regions (HAR) or human-gained (HGE) or lossed enhancer (HLE) regions (*48*). However, no enrichments for SA or CT were identified for HAR, HGE and HLE genes (Fig. 5D-F). These results expand on the notion that human-specific genes are of relevance to ASD by showing that the normative genomic mechanisms associated to SA are also genes of importance for human-specific evolution. Given that the SA LV1 relationship is absent in ASD Poor, this suggests that the loss of such normative associations may allow for early SA expansion and possibly early brain overgrowth for ASD Poor.

Next, we asked whether SA and CT non-zero modules were relevant for known autism-associated genomic mechanisms. SA LV1 and CT LV1 non-zero or zero modules are not enriched for rare de novo protein truncating variants (*49*) or other genes that are annotated as autism-associated in SFARI Gene (*50*). However, CT LV2 non-zero modules were enriched for SFARI ASD genes (Table S1). Thus, at the level of ASD-risk gene mutations, CT LV2 was the only feature showing enrichments with non-zero modules. This could be compatible with the nature of CT LV2 being mostly specific to the ASD Poor subtype.

At the level of genes with evidence of ASD-dysregulated expression from post-mortem cortical tissue, we find that both CT LV1 and LV2 non-zero modules were enriched for ASD upregulated genes (*51*). In contrast, genes from cortically downregulated co-expression modules (*19*) were highly enriched with genes from SA LV1 non-zero modules (Table S1). This result shows an interesting contrast between CT and genes that show upregulated expression versus SA and genes that show downregulated expression in ASD.

Non-zero modules from SA LV1, CT LV1, and CT LV2 are also enriched for co-expression modules that are highly transcriptionally active during prenatal periods and which contain many high-penetrance ASD-related mutations (Fig. 6A-C; Table S1). This is compatible with the idea that broadly expressed genes can interact and impact key ASD-risk genes, particularly in prenatal periods (*9, 10*). Downstream targets of highly penetrant genes like *FMR1* and *CHD8* were also enriched in non-zero modules from SA LV1, CT LV1, and CT LV2. However, not all of these enrichments are specific to autism-associated genes. Genes differentially expressed in schizophrenia (*51*) were also significantly enriched in non-zero modules across SA LV1, CT LV1, and CT LV2.

**Fig. 6:**
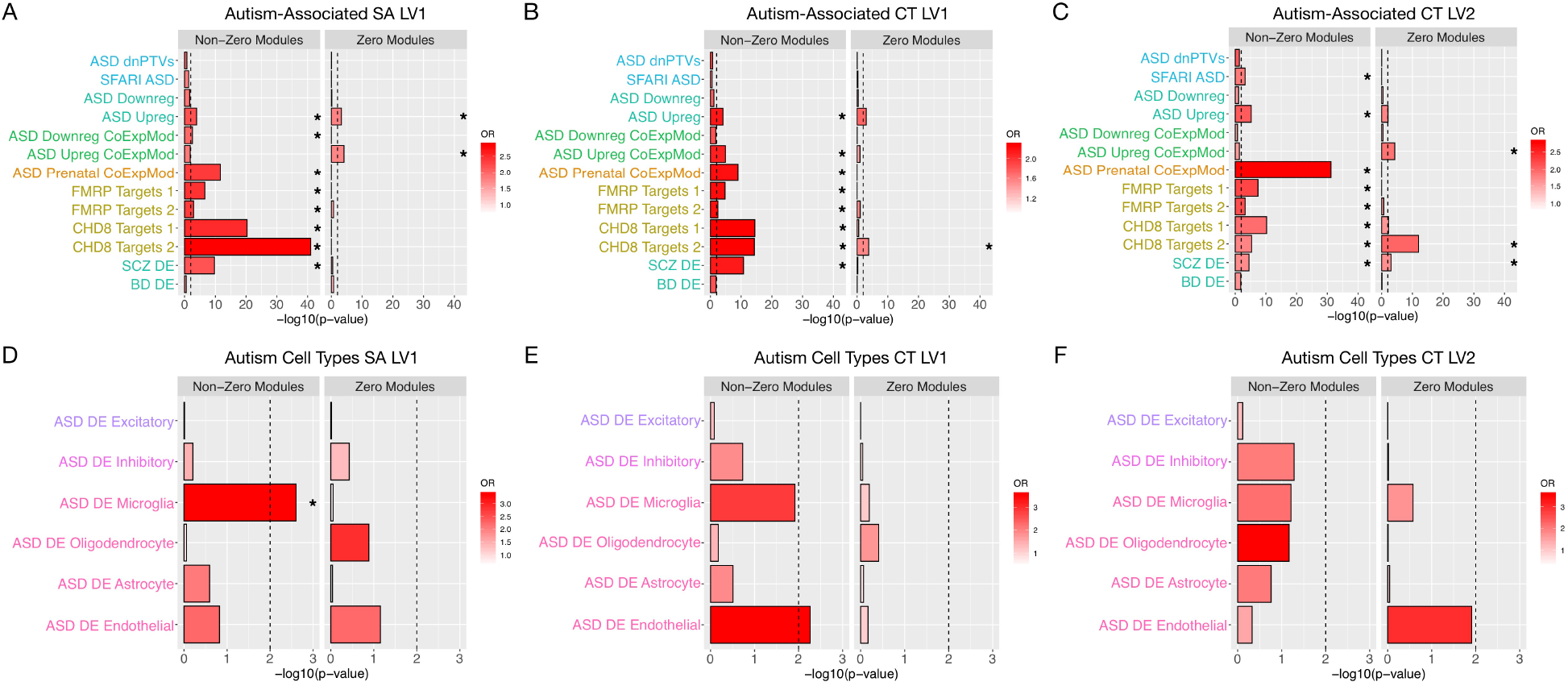
Enrichment between PLS non-zero modules and autism-associated genes. Panels A-C indicate enrichments between different autism-associated gene lists and non-zero and zero modules for SA LV1 (A), CT LV1 (B), AND CT LV2 (C). Panels D-F indicate enrichments between differentially expressed genes in specific cell types in autism and non-zero and zero modules for SA LV1 (D), CT LV1 (E), and CT LV2 (F). Asterisks marks enrichments at FDR q<0.01. Abbreviations: DE, differentially expressed; OR, enrichment odds ratio; SA, surface area; CT, cortical thickness; LV, latent variable pair; dnPTVs, de novo protein truncating variants.

Finally, we examined enrichments with cell type specific differentially expressed genes in autism (*52*). Here we found that only SA LV1 non-zero modules are enriched for differentially expressed genes in microglia cells (Fig. 6D). No other comparisons for DE cell types were statistically significant. See Fig. 6 and Table S1 for a summary of autism-associated enrichments. The fact that non-zero modules are devoid of enrichments in most DE genes from specific cell types is compatible with the notion that these genes are of primary relevance for early prenatal periods and will not be a highly discoverable DE signal in post-mortem ASD tissue.

To aid future work examining specific genes of interest, we focused on identifying high-confidence ASD-risk genes (annotated as the ‘high-confidence’ category 1 list in SFARI Gene) that are also SA-and prenatally-relevant progenitor and A-P patterning genes (i.e. the intersection of SFARI ASD, SA non-zero modules, PC1 A-P genes, prenatal progenitor cell types, and ASD prenatal co-expression modules). *SON* and *BAZ2B* were identified and these genes play roles in splicing, cell cycle, transcriptional regulation, and chromatin remodeling. For CT LV1 genes, we next searched for high-confidence ASD-risk genes that were also prenatally-relevant excitatory and D-V patterning genes (i.e. the intersection of SFARI ASD, CT LV1 non-zero modules, PC2 D-V genes, prenatal excitatory cell types, and ASD prenatal co-expression modules). Here we find ASD ‘high-confidence’ genes of *ATRX*, *AUTS2*, and *BCL11A*. In a similar search within CT LV2 non-zero modules of prenatal relevance to excitatory neurons and D-V patterning, we identified *ATRX*, *AUTS2*, *BCL11A*, *CACNA1E*, and *MEIS2* as high-confidence ASD-risk genes. A common theme of all these CT-relevant genes is their role in chromatin modification and remodeling (with the exception of *CACNA1E*) and their links to syndromes causing intellectual disability. Additionally, with the exceptions of *BCL11A* and *CACNA1E*, all SA- and CT-relevant high-confidence genes listed here fall into the broadly expressed gene list, highlighting the importance of these high-impact genes in ASD biology (*9*).

These findings represent a significant enhancement to the mechanistic and clinical precision of our understanding of the early brain basis behind the autisms (*1*). Along with our prior work (*6, 7*), this work showcases that the ASD Poor subtype indeed a distinct subtype with multiscale differentiation across development, behavior, and underlying neural systems. While prior work showed a biological distinction in this subtype with fMRI (*6, 7*), this work shows that the biology is also distinct when examining structural neural phenotypes like SA and CT. While on-average early brain overgrowth is one of the most robust findings in the literature on neurodevelopment in ASD (*4, 31*), this work shows that the effect is driven by a subtype of ASD toddlers with poor early language development and outcome.

This work also uncovers an altogether new discovery behind how cortical SA and CT phenotypes are atypically genomically patterned in the ASD Poor subtype. Genomic patterning of SA and CT occurs along A-P and D-V gradients, thus enabling the development of cortical areal and circuit identities (*11*, *13*, *25*–*27*). This A-P and D-V genomic patterning of SA and CT is intact in TD and ASD Good, but absent in ASD Poor (e.g., Fig. 2D). Atypical genomic patterning of the cortex in ASD Poor could be the key neural explanation behind why these individuals have much more pervasive and more severe behavioral difficulties and poor outcomes. Prior work has shown that molecular identity defined by gene expression affects cell type specific neurophysiological response (*53*). Thus, without intact genomic patterning of the cortex it may be that development of regional or circuit level identities may be perturbed in ASD Poor and this may help explain the phenotypic difficulties in complex information processing in domains like language and social communication.

The results also shed insight into the developmental and mechanistic origins at the root of the ASD Poor subtype. Evidence suggests that these SA and CT-relevant genes are the same genes responsible in early prenatal periods for establishing these A-P and D-V gene expression gradients across the cortex. The genes responsible for this atypical prenatal genomic patterning are massive in scale, encompassing a large majority of the genes examined. This result is compatible with ideas from the omnigenic model of complex traits (*8*). The omnigenic model also proposed that broadly expressed genes should have large impact on complex traits encompassed by neuropsychiatric phenotypes and diagnoses. Indeed, broadly expressed genes manifest in this and other studies (*6*) and are a key class of ASD genetic risk that operates at early prenatal timepoints (*9, 10*). In future work it will be important to explore how genomic patterning of the cortex can affect other ASD subtypes. Additionally, it will be important for future work to investigate how omnigenic and broadly expressed genes such as those identified here may play roles in other atypical multiscale phenomena in ASD.

## Acknowledgments

We thank all participants and their families for participating in this study.

## Funding

This project has received funding from the European Research Council (ERC) under the European Union's Horizon 2020 research and innovation programme under grant agreement No 755816 (ERC Starting Grant to MVL). This work was also supported by the following grants to EC, KP, LE, and NEL - NIMH R01-MH080134 (KP), NIMH R01-MH104446 (KP), NFAR grant (KP), NIMH Autism Center of Excellence grant P50-MH081755 (EC, KP), NIMH R01-MH036840 (EC), NIMH R01-MH110558 (EC, NEL), NIMH U01-MH108898 (EC), NIDCD R01-DC016385 (EC, KP, LE, MVL), CDMRP AR130409 (EC), and the Simons Foundation 176540 (EC). KC was supported by the Utah Stimulating Access to Research in Residency Transition Scholar (StARRTS) under Award Number 1R38HL143605-01.

## Author contributions

Conceptualization: MVL, EC, KP, LE TP. Methodology: MVL, TP, VHG, NEL, EC, KP, LE. Software: MVL, DJH, AMD, VHG. Formal analysis: MVL, VHG, DJH, TP, JS, RAIB, NB. Investigation: LE, KC, CCB, LL, KP, EC. Data curation: EC, KP, LE, TP, VHG, MVL. Writing -original draft preparation: MVL, EC. Writing - review and editing: MVL, EC, KP, LE, JS RAIB, NB, KC, NEL, VHG, DJH, AMD. Visualization: MVL. Supervision: EC, KP, LE, NEL. Project administration: EC, KP, LE, NEL. Funding acquisition: MVL, EC, KP, LE, NEL.

## Competing interests

None of the authors have any biomedical financial interests or potential conflicts of interest to report.

## Data and materials availability

Analysis code is available at https://github.com/IIT-LAND/genomic_cortical_patterning_autisms. Data are publicly available from the NIH National Database for Autism Research (NDAR). Raw and normalized blood gene expression data are also deposited in Gene Expression Omnibus (GSE42133; GSE111175). RNA-seq data from the Development PsychENCODE dataset can be found here: http://development.psychencode.org. GTEx data can be found here: https://www.gtexportal.org. Microarray data from the songbird vocal learning model can be found in Gene Expression Omnibus (GSE34819).

## Supplementary Materials

### Materials and Methods

#### Participants

This study was approved by the Institutional Review Board at University of California, San Diego. Parents provided written informed consent according to the Declaration of Helsinki and were paid for their participation. Identical to the approach used in our earlier studies (*6*, *7*, *22*, *54*–*59*) toddlers were recruited through two mechanisms: community referrals (e.g., website) or a general population-based screening method called the 1-Year Well-Baby Check-Up Approach (*60*) that allowed for the prospective study of ASD beginning at 12 months based on a toddler’s failure of the CSBS-DP Infant-Toddler Checklist (*61, 62*). All toddlers were tracked from an intake assessment around 12 months and followed roughly every 12 months until 3–4 years of age. All toddlers, including normal control subjects, participated in a series of tests collected longitudinally across all visits, including the Autism Diagnostic Observation Schedule (ADOS; Module T, 1, or 2) (*63*), the Mullen Scales of Early Learning (*64*), and the Vineland Adaptive Behavior Scales (*65*). All testing occurred at the University of California, San Diego Autism Center of Excellence (ACE). No randomization procedures were implemented as part of the data collection process. Data collection and analyses were not performed blind to the conditions of the experiment.

Stratification of ASD Poor versus ASD Good was made on the basis of Mullen EL and RL T-scores. An ASD toddler was classified as ASD Poor if both Mullen EL and RL T-scores at the final outcome assessment was below 1 standard deviation of the T-score norm of 50 (i.e. T<40). ASD Good labels were made if the toddler had either Mullen EL or RL T-scores within 1 standard deviation or above the normative T-score of 50 (i.e. T ≥ 40). A total of n=123 toddlers had T1 structural MRI and gene expression data available. From these 123 toddlers, n=76 ASD individuals were examined and were split into the 2 language outcome subtypes - ASD Poor n=38 (32 male, 6 female; mean age at MRI scan = 29.01 months, SD at fMRI scan = 7.22, range = 12-50 months), ASD Good n=38 (28 male, 10 female; mean age at fMRI scan = 29.02 months, SD at fMRI scan = 9.55, range = 14-46 months) and TD n=47 (25 male, 22 female; mean age at fMRI scan = 25.91 months, SD at fMRI scan = 10.44, range = 13-46 months). ASD subtypes and TD did not statistically differ in age at the time of scanning (*F(2,120)* = 1.62, *p* = 0.20). For more demographic and phenotypic information, please see Table S2.

#### Blood Sample Collection, RNA extraction, quality control and samples preparation

Four to six milliliters of blood was collected into EDTA-coated tubes from toddlers on visits when they had no fever, cold, flu, infections or other illnesses, or use of medications for illnesses 72 hours prior blood draw. Blood samples were passed over a LeukoLOCK™ filter (Ambion, Austin, TX, USA) to capture and stabilize leukocytes and immediately placed in a −20°C freezer. Total RNA was extracted following standard procedures and manufacturer’s instructions (Ambion, Austin, TX, USA). LeukoLOCK disks (Ambion Cat #1933) were freed from RNA-later and Tri-reagent (Ambion Cat #9738) was used to flush out the captured lymphocyte and lyse the cells. RNA was subsequently precipitated with ethanol and purified though washing and cartridge-based steps. The quality of mRNA samples was quantified by the RNA Integrity Number (RIN), values of 7.0 or greater were considered acceptable (*66*), and all processed RNA samples passed RIN quality control. Quantification of RNA was performed using Nanodrop (Thermo Scientific, Wilmington, DE, USA). Samples were prepped in 96-well plates at the concentration of 25 ng/µl.

#### Gene expression and data processing

RNA was assayed at Scripps Genomic Medicine (La Jolla, CA, USA) for labeling, hybridization, and scanning using the Illumina BeadChips pipeline (Illumina, San Diego, CA, USA) per the manufacturer’s instruction. All arrays were scanned with the Illumina BeadArray Reader and read into Illumina GenomeStudio software (version 1.1.1). Raw data was exported from Illumina GenomeStudio, and data pre-processing was performed using the lumi package (*67*) for R (http://www.R-project.org) and Bioconductor (https://www.bioconductor.org) (*68*). Raw and normalized data are part of larger sets deposited in the Gene Expression Omnibus database (GSE42133; GSE111175).

A larger primary dataset of blood leukocyte gene expression was available from 383 samples from 314 toddlers with the age range of 1-to-4 years old. The samples were assayed using the Illumina microarray platform on three batches. The datasets were combined by matching the Illumina Probe ID and probe nucleotide sequences. The final set included a total of 20,194 gene probes. Quality control analysis was performed to identify and remove 23 outlier samples from the dataset. Samples were marked as outlier if they showed low signal intensity (average signal two standard deviations lower than the overall mean), deviant pairwise correlations, deviant cumulative distributions, deviant multi-dimensional scaling plots, or poor hierarchical clustering, as described elsewhere (*55*). The high-quality dataset included 360 samples from 299 toddlers. High reproducibility was observed across technical replicates (mean Spearman correlation of 0.97 and median of 0.98). Thus, we randomly removed one of each of two technical replicates from the primary dataset. From the subjects in the larger primary dataset, n=123 also had MRI data and thus a total of n=105 from the Illumina HT12 platform along with n=18 from the Illumina WG6 platform were used in this study. Batch was not asymmetrically distributed across one subgroup more than another, as chi-square analyses on the contingency table between subgroup and batch show no effect (*χ*^*2*^*(4)* = 0.84, *p* = 0.93). ASD subtypes and TD toddlers also did not statistically differ in age at the time of blood sampling (*F(2,120)* = 1.27, *p* = 0.28). The 20,194 probes were then collapsed to 14,426 genes based on picking the probe with maximal mean expression across samples. Data were quantile normalized and then adjusted for batch effects, sex, and RIN. This batch, sex, and RIN adjusted data were utilized in all further downstream analyses. We also checked for differences in proportion estimates of different leukocyte cell types (i.e. neutrophils, B cells, T cells, NK cells, and monocytes) using the CellCODE deconvolution method (*69*), but found no evidence of differences across groups for any cell type (see Table S3).

#### Weighted Gene Co-Expression Network Analysis

We reduced the number of features in the gene expression dataset from 14,426 genes down to 21 modules of tightly co-expressed genes. This data reduction step was achieved using weighted gene co-expression network analysis (WGCNA), implemented within the WGCNA library in R (*34*). Correlation matrices estimated with the robust correlation measure of biweight midcorrelation were computed and then converted into adjacency matrices that retain the sign of the correlation. These adjacency matrices were then raised to a soft power of 16 (Fig. S2). This soft power was chosen by finding the first soft power where a measure of R^2^ scale-free topology model fit saturates. The soft power thresholded adjacency matrix was then converted into a topological overlap matrix (TOM) and then a TOM dissimilarity matrix (e.g., 1-TOM). The TOM dissimilarity matrix was then input into agglomerative hierarchical clustering using the average linkage method. Gene modules were defined from the resulting clustering tree, and branches were cut using a hybrid dynamic tree cutting algorithm (deepSplit parameter = 4) (Fig. S2). Modules were merged at a cut height of 0.2, and the minimum module size was set to 100. Only genes with a module membership of r > 0.2 were retained within modules. For each gene module, a summary measure called the module eigengene (ME) was computed as the first principal component of the scaled (standardized) module expression profiles. We also computed module membership for each gene and module. Module membership indicates the correlation between each gene and the module eigengene (see Table S4). Genes that could not be clustered into any specific module are left within the M0 module, and this module was not considered in any further analyses. Further WGCNA analyses were run separately within each group in order to check for preservation of detected modules across groups at a soft power threshold of 16. These analyses all indicated high levels of preservation (Zsummary>10) (*70*) for all detected modules for each pairwise group comparison (Fig. S3).

#### MRI Data Acquisition and Analyses

Imaging data were collected on a 1.5 Tesla General Electric MRI scanner during natural sleep at night; no sedation was used. Structural MRI data was collected with a T1-weighted IR-FSPGR sagittal protocol (TE = 2.8 ms, TR = 6.5 ms, flip angle = 12 degrees, bandwidth = 31.25 kHz, FOV = 24 cm, slice thickness = 1.2 mm). Cortical surface reconstruction was performed using FreeSurfer v5.3 (http://surfer.nmr.mgh.harvard.edu/) (*71–73*), which uses routinely acquired T1-weighted MRI volumes (*74*), includes tools for estimation of brain morphometry measures such as cortical thickness and surface area (*75, 76*), and enables inter-subject alignment via nonlinear, surface-based registration to an average brain, driven by cortical folding patterns (*77*). FreeSurfer has been validated for use in children (*78*) and used successfully in large pediatric studies (*79, 80*). Total cortical volume, surface area (SA) and mean cortical thickness (CT) were computed based on the Desikan-Killiany parcellation. Regional SA and CT values were computed from a 12-region parcellation reported by Chen and colleagues (*26, 27*) based on genetic similarity in monozygotic twins. This parcellation scheme, known as GCLUST, is highly relevant for our purposes here, since the parcellations are based on genetic patterning. Thus, GCLUST should help increase statistical power while also minimizing multiple comparisons. The GCLUST parcellation is also important as it can be used to leverage information about genetic similarity gradients (e.g., rank ordering of regions by fuzzy clustering) in further analyses. The 2-cluster anterior-posterior (A-P) or dorsal-ventral (D-V) partitions discovered by Chen and colleagues (*26, 27*) are also relevant in further analyses for A-P and D-V gradient questions. For all 12 regions of the SA and CT GCLUST parcellation, global effects were controlled for by dividing SA values by the mean SA, and for CT we subtracted mean CT from each region, as was done in prior papers using this parcellation scheme (*26, 27*).

#### MRI-Gene Expression Association Analysis

To assess multivariate MRI-gene expression relationships we used partial least squares (PLS) analysis (*81, 82*). PLS is widely used in the neuroimaging literature, particularly when explaining multivariate neural responses in terms of multivariate behavioral patterns of variation or a design matrix. Given that the current dataset is massively multivariate both in terms of MRI and gene expression datasets, we used PLS to elucidate how variation in SA or CT covaries with gene expression as measured by module eigengene values of co-expression modules. PLS allows for identifying such relationships by finding latent MRI-gene expression variable pairs (LV) that maximally explain covariation in the dataset and which are uncorrelated with other MRI-gene expression LV pairs. The strength of such covariation is denoted by the singular value (d) for each brain-gene expression LV, and hypothesis tests are made via using permutation tests on the singular values. Furthermore, identifying brain regions that most strongly contribute to each LV pair is acheived via bootstrapping, whereby a brain bootstrap ratio (BSR) is created for each region, and represents the reliability of that region for contributing strongly to the LV pattern identified. The brain BSR is roughly equivalent to a Z-statistic and can be used to threshold data to find voxels that reliably contribute to an LV pair.

The PLS analyses reported here were implemented within the plsgui MATLAB toolbox (www.rotman-baycrest.on.ca/pls/). Here we ran 2 separate PLS analyses - one on SA and another on CT. Neuroimaging data entered into the PLS analyses come from the 12 region GCLUST parcellations for SA and CT. Because the TD group differed in the proportion and males versus females compared to the ASD groups, we used a linear model to remove the effect of sex from the SA and CT data. This SA and CT data with the sex effect removed was input into the PLS analysis. For gene expression data, we input module eigengene values for all 21 co-expression modules. For statistical inference on identified MRI-gene expression LV pairs, a permutation test was run with 10,000 permutations. To identify reliably contributing regions for MRI-gene expression LVs and to compute 95% confidence intervals (CIs) on MRI-gene expression correlations, bootstrapping was used with 10,000 resamples. Gene co-expression modules whereby 95% CIs do not encompass 0 are denoted as ‘non-zero’ association modules. All other modules where 95% CIs include 0 are denoted as ‘zero’ modules.

From the PLS results we tested whether groups show similar correlation patterns across modules. To test this question, we computed Pearson correlations on the PLS correlation values for all pairwise group comparisons. Groups with similar PLS correlations will show statistically significant correlations. We also used the brain bootstrap ratios (BSR) from the PLS analysis to identify whether BSRs covary along the genetic similarity gradients and A-P and D-V partitions discovered by Chen and colleagues (*26, 27*). Pearson correlations were used to identify correlations with genetic similarity gradients, while independent-samples t-tests were used to compare A-P and D-V partitions.

#### Gene Set Enrichment Analyses

We analyze enrichment between genes from PLS non-zero and zero modules and a host of other gene lists defined by a variety of criteria (see below for details). For these gene set enrichment analyses, we utilized custom R code written by MVL (https://github.com/mvlombardo/utils/blob/master/genelistOverlap.R) that computes hypergeometric p-values and enrichment odds ratios. The background pool for these enrichment tests was always set to 14,426. After all enrichment tests were computed, results are interpreted only if the enrichment was statistically significant after FDR correction for multiple comparisons at a threshold of FDR q<0.01.

#### Prenatal Gene Expression Gradients and Cell Types

To assess gradients in prenatal gene expression we utilized RNA-seq data from the Development PsychENCODE dataset (http://development.psychencode.org) (*14*). The data utilized was already preprocessed as described by Li and colleagues (*14*) (e.g., normalized, batch effects removed) and summarized to RPKM. Sample data from all 12 available cortical regions from 12-22 weeks post-conception were utilized in order to capture the midgestational window of interest. Before running the analysis we removed low expressing genes with log2(RPKM) below 2. The primary analysis to identify expression gradients was an adjustment-for-confounds principal components analysis (AC-PCA) (*37*) which allowed for adjustment due to repeat measurements from the same donor across sampled brain regions. Rank ordering of regions by A-P and D-V axes were utilized to statistically confirm that PC1 and PC2 components follow A-P and D-V gradients. Subsets of the most important genes for the top two principal components were identified with a sparse AC-PCA analysis, whereby the sparsity parameter, c2, was selected based on a grid search with 10-fold cross validation. These PC1 and PC2 gene sets were used in enrichment tests with PLS non-zero or zero modules.

We also examined enrichments between PLS non-zero and zero modules and prenatal cell types identified from single cell RNA-seq on midgestational prenatal brain tissue (*42*). These cell types included several classes of progenitor cells (ventricular radial glia, vRG; outer radial glia, oRG; cycling progenitors (S phase), PgS; cycling progenitors (G2/M phase), PgG2M; intermediate progenitors, IP), excitatory neurons (migrating excitatory, ExN; maturing excitatory, ExM; maturing excitatory upper enriched, ExM-U; excitatory deep layer 1, ExDp1; excitatory deep layer 2, ExDp2), inhibitory neurons (interneuron CGE, InCGE; interneuron MGE, InMGE), and other non-neuronal cell types (oligodendrocyte precursors, OPC; pericytes, Per; endothelial cells, End; microglia, Mic).

#### Tissue-specific enrichments

To better understand how genes expressed in blood leukocytes could be brain-relevant we annotated gene co-expression modules based on enrichments in genes known from expression across multiple tissues to be either broadly expressed or brain-specific. Both of these categories contain genes that are expressed in cortical tissue, but differ in the pattern of expression across other non-neuronal tissues. To define these lists we downloaded transcript per million (TPM) normalized gene expression from 10,259 samples across 26 tissues from the GTEx dataset (https://www.gtexportal.org) (*83, 84*). In addition to brain and nerve tissue, the dataset included transcriptome data from 24 non-neuronal tissues, including: Adipose, Adrenal Gland, Blood Vessel, Breast, Blood, Skin, Colon, Esophagus, Heart, Liver, Lung, Salivary Gland, Muscle, Ovary, Pancreas, Pituitary, Prostate, Small Intestine, Spleen, Stomach, Testis, Thyroid, Uterus, and Vagina (Table S2). We next defined a gene expressed in a tissue if it met two criteria. First, the gene TPM expression level was ≥ 3 in at least half of the samples from the tissue. Second, the median expression of the gene was equal or larger than its 25-percentile expression in GTEx cortex samples. The second criterion was included to account for the differences in the base expression level of the genes and their dosage dependent translation and function. Broadly-expressed genes were defined as genes that were expressed in ≥ 50% of non-neuronal tissues (i.e., tissues other than brain and nerve). The broadly-expressed and brain-specific genes included genes that were expressed in the adult cortex based on GTEx dataset.

#### Vocal learning enrichments

To test for enrichment between PLS non-zero modules and gene sets of functional relevance for language processes, we examined genes that are differentially expressed in a song bird vocal learning model. Song birds are often used as animal models relevant for the vocal learning component of language (*44*, *85*, *86*). We investigated enrichments with differentially expressed genes taken from a microarray dataset of Area X of song birds (*44*). To identify differentially expressed (DE) genes between singing versus non-singing birds, we re-analyzed this dataset (GEO Accession ID: GSE34819) using limma (*87*), and DE genes were identified if they passed Storey FDR q<0.05 (*88*). These DE genes were also used for enrichment tests in our prior work examining gene expression relationships with language-relevant functional neural phenotypes measured with fMRI (*6*).

#### Human-specific enrichments

Given the uniquely human nature of language, we also tested hypotheses regarding enrichments with genes that are transcriptionally different in the cortical tissue between humans and other non-human primates across prenatal, early postnatal and adult periods (*47*). In addition, we also examined enrichments with genes linked to human accelerated regions (HAR), human-gained enhancers (HGE) in prenatal and adult tissue, and human-lossed enhancers (HLE) (*48*).

#### Autism-associated enrichments

Ample evidence suggests that prenatal periods are critical for ASD (*4*, *9*, *89*–*91*). To test enrichment with prenatal ASD-associated co-expression modules, we utilized co-expression modules from a study that analyzed the Allen Institute BrainSpan dataset (*92*). Parikshak and colleagues analyzed only cortical regions from BrainSpan and identified M2 and M3 as prenatally active and enriched for rare protein truncating variants with high penetrance for ASD (*89*). We also tested enrichments with gene lists known to be associated with ASD, either from genetic evidence or evidence from cortical transcriptomic dysregulation. In particular, we examined a list of 102 rare de novo protein-truncating variants (dnPTV) associated with ASD (*49*), genes listed as ASD-associated in SFARI Gene (https://gene.sfari.org) in categories S, 1, 2, and 3 (downloaded on July 16, 2020) (*50*), and DE genes and cortical co-expression modules measured from ASD post-mortem frontal and temporal cortex tissue (*19, 51*). To contrast ASD DE genes to genes that are DE in other psychiatric diagnoses that are genetically correlated with autism, we also use DE genes in schizophrenia (SCZ DE) and bipolar disorder (BD DE) from the same study that identified ASD DE genes (*51*). To go beyond DE genes identified in bulk tissue samples, we also examined ASD DE genes identified in specific cell types - particularly, excitatory (ASD Excitatory) and inhibitory (ASD Inhibitory) neurons, microglia (ASD Microglia), astrocytes (ASD Astrocyte), oligodendrocytes (Oligodendrocyte), and endothelial (ASD Endothelial) cells (*52*). Finally, we also tested for enrichments with known downstream targets of highly penetrant mutations known to be associated with ASD – FMRP and CHD8. For each, we had lists of downstream targets for two independent studies (*93–96*), where the overlap for FMRP targets was 3.71% and 27.61% for CHD8 targets.

## Figures S1-S3

**Fig. S1:**
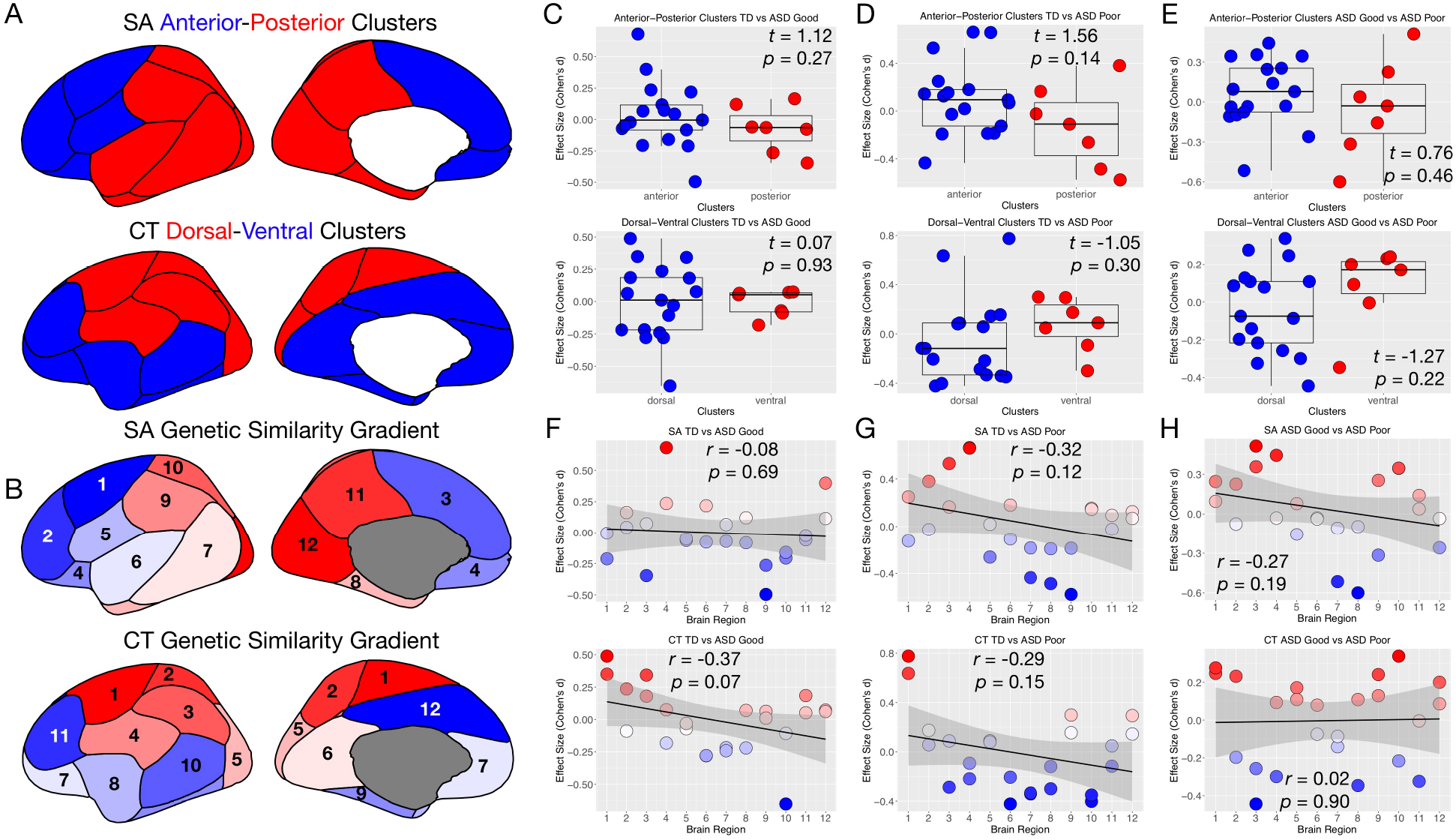
Lack of A-P and D-V gradients in effect size of group comparisons. Panel A shows a depiction of the A-P and D-V partitions defined by Chen and colleagues (26, 27). Panel B shows how the genetic similarity gradient defined by Chen and colleagues (26, 27) manifests via numbered ordering of brain regions along that gradient. Panels C-E show standardized effect size (Cohen’s d) for group comparisons of TD vs ASD Good (C), TD vs ASD Poor (D), and ASD Good vs ASD Poor (E). The top row of panels C-E are for the A-P partition, while the bottom row in each panel is for the D-V partition. Panels F-H show scatterplots of standardized effect size by genetic similarity gradient for each group comparison of TD vs ASD Good (F), TD vs ASD Poor (G), and ASD Good vs ASD Poor (H). Comparisons for SA are shown in the top of each panel in F-H, while CT is shown at the bottom.

**Fig. S2:**
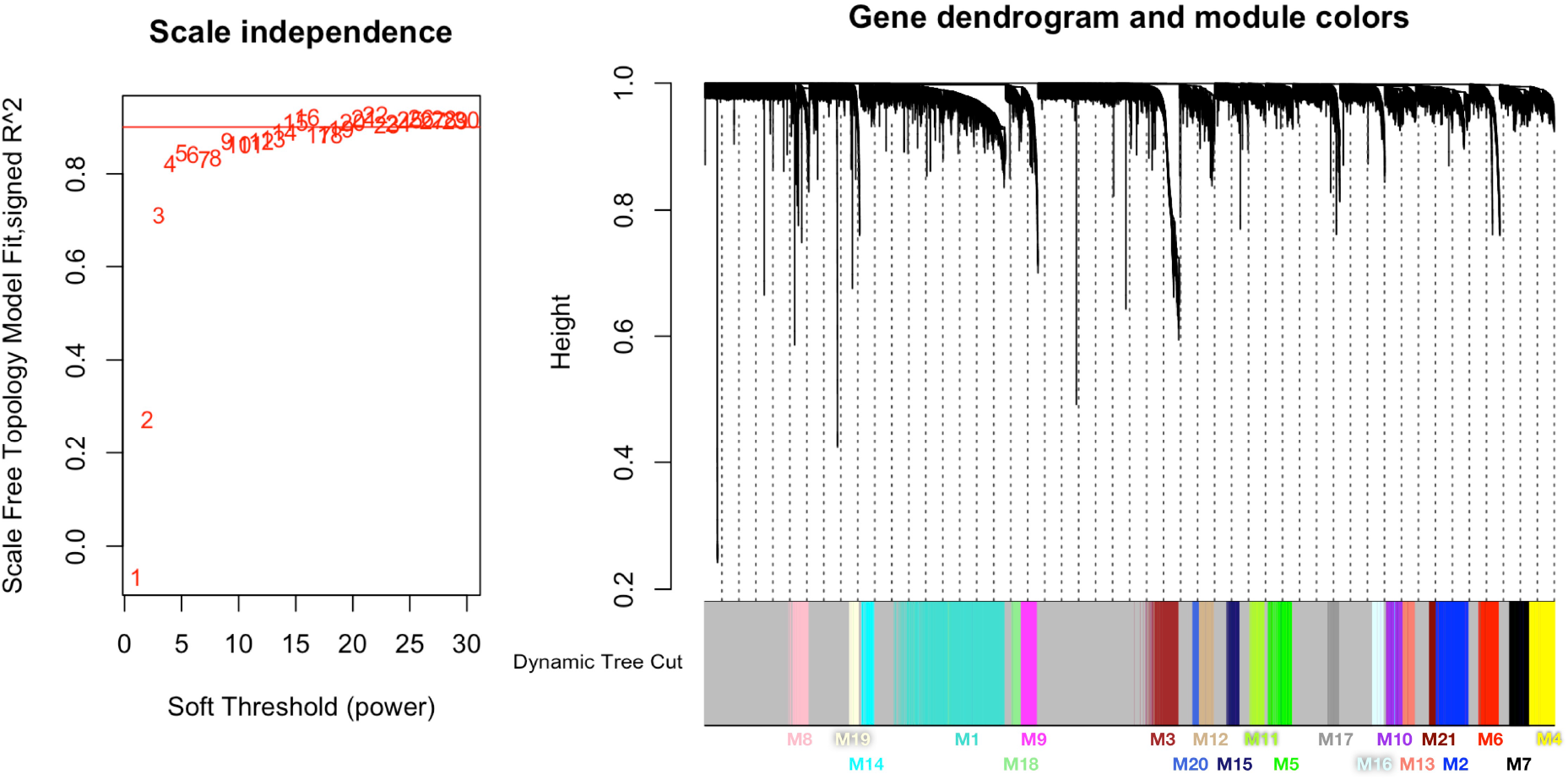
Soft power and TOM dendrogram from WGCNA analysis. On the left of this figure we show the soft power plot for the main WGCNA analysis including data from all groups. A horizontal red line depicts soft power topology model fit R^2^ of 0.9, where the chosen soft power of 16 is located. On the right of this figure is the TOM dendrogram with modules labeled at the bottom.

**Fig. S3:**
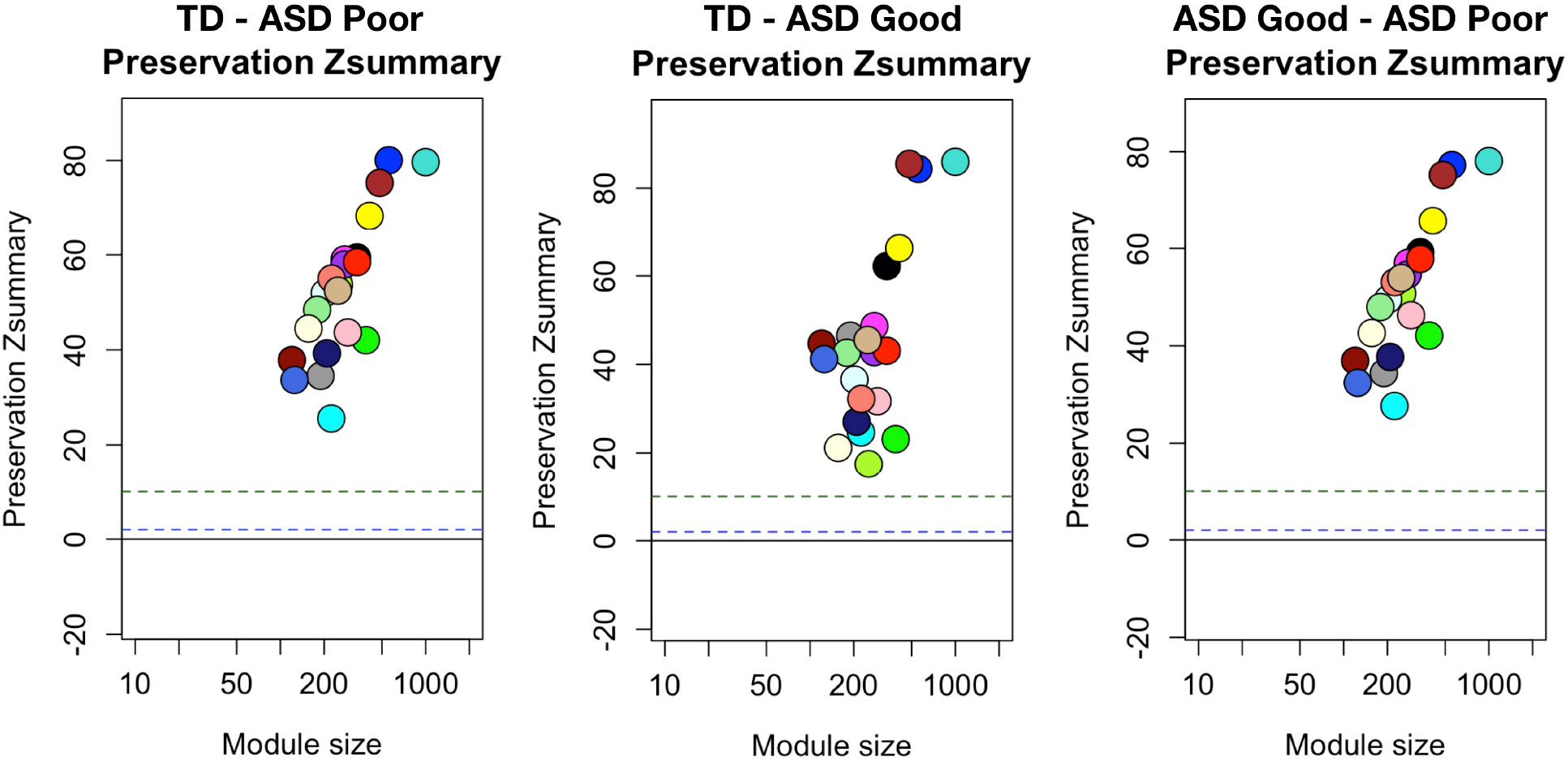
Module preservation when WGCNA analysis is run separately on each group. This figure shows the module preservation Zsummary statistic for WGCNA analyses run separately on each group in order to show that networks are highly preserved (Zsummary>10) across groups.

## Tables S1-S4

Table S1: Table annotating overlap of each gene with gene sets used in enrichment analyses.

Table S2: Summary of clinical and demographic variables.

Table S3: ANOVA stats from CellCODE deconvolution of leukocyte cell types.

Table S4: WGCNA module assignments for each gene and module membership scores.

## Notes

### Competing Interest Statement

The authors have declared no competing interest.

